# Chronic optogenetic activation of hippocampal pyramidal neurons replicates the proteome footprint of Alzheimer’s disease-like pathology

**DOI:** 10.1101/2023.09.12.557365

**Authors:** Iason Keramidis, Martina Samiotaki, Romain Sansonetti, Johanna Alonso, Katerina Papanikolopoulou, Yves De Koninck

**Affiliations:** CERVO Brain Research Centre, Quebec Mental Health Institute, Québec, QC, Canada; Department of Psychiatry & Neuroscience, Université Laval, Québec, QC, Canada; Institute for bio-Innovation, Biomedical Sciences Research Center Alexander Fleming, Vari, Greece; Institute for Fundamental Biomedical Research, Biomedical Sciences Research Center Alexander Fleming, Vari, Greece

## Abstract

Neuronal overexcitability can elicit synaptic changes, leading to neuronal hyperactivity and abnormal neural circuit processing. Such network disruption impairs neuronal function and survival, initiating neurodegeneration and Alzheimer’s disease (AD). Yet, the sequence of synaptic changes resulting from excessive neuronal activity remains elusive. We employed optogenetics to induce sustained neuronal hyperactivity in the hippocampi of wild-type and AD-like 5xFAD mice. Surprisingly, after a month of daily optogenetic stimulation, the proteomic profiles of photoactivated wild-type and 5xFAD mice exhibited remarkable similarity. Proteins involved in translation, protein transport, autophagy, and notably in the AD pathology were upregulated in wild-type mice. Conversely, both glutamatergic and GABAergic synaptic proteins were downregulated. These hippocampal proteomic and signaling alterations in wild-type mice resulted in spatial memory loss and augmented Αβ42 secretion. Collectively, these findings indicate that sustained neuronal hyperactivity alone replicates proteome changes seen in AD-linked mutant mice. Therefore, prolonged neuronal hyperactivity may contribute to synaptic transmission disruption, memory deficits and the neurodegenerative process associated with AD.

## Introduction

Aberrant neuronal activity can impair synapse formation and strength resulting in disruption of synaptic plasticity. In turn, altered synaptic plasticity could reorganize synaptic connections ensuing runaway excitation or quiescence and eventually modify local neuronal circuits (*1*). For neural circuits to maintain their characteristic firing patterns, neurons within the network undergo homeostatic changes at the cellular level, regulating synthesis and degradation of key synaptic proteins (*2*, *3*). Alterations perturbing these regulations have been linked with the onset of various brain disorders such as epilepsy and autism (*4*).

Neuronal hyperexcitability manifests early in Alzheimer’s disease (AD) (*5–7*), leading to cortical and hippocampal hyperactivity (*8*, *9*) and under certain conditions to epileptiform activity and seizures in rodents (*10*, *11*) and humans (*12*). Both amyloid-β (Aβ) and Tau have been found to induce neuronal hyperactivity through distinct cellular mechanisms (*11*, *13–16*). Concurrently, neuronal hyperactivity can increase Αβ secretion (*17*, *18*) and promote pathological phosphorylation of Tau (*19*). These observations suggest that Αβ and Tau can initiate a vicious cycle of neuronal hyperactivity in AD underlying neurodegeneration in disease. However, it remains unknown whether neuronal hyperactivity alone can induce pathology mimicking neurodegeneration in AD.

Optogenetics has emerged as a powerful method to control the activity of genetically defined neurons allowing to study the function of selective brain circuits in physiological but also pathological conditions (*20–22*). Stabilized step function opsins (SSFOs) allow also for long-lasting activation of neural circuits with just a brief light stimulation (*23*). Such chronic optogenetic activation is necessary to model neurodegenerative disorders like AD for which neuronal activity is altered for long periods of time. We adopted a similar approach to generate a model of chronic neuronal hyperactivity in the hippocampus of wild-type (WT) and a transgenic mouse line carrying AD-linked mutations and demonstrated that evoked neuronal hyperactivity can disrupt synaptic signalling and facilitate an activity-driven neuropathology similar to AD.

## Results

### Optogenetic activation of the hippocampus in wild-type and 5xFAD mice using SSFO

To generate a model of chronic neuronal hyperactivity in the rodent hippocampus we virally transduced the SSFO-mCherry or tdTomato unilaterally under the CaM kinase II promoter (CaMKIIa) into the CA1 of WT or 5xFAD (*24*) male mice (Fig. 1A, B). We then inserted an optic fiber cannula above the CA1 and stimulated the infected neurons for 4 weeks by blue light (Fig. 1A). To validate whether light stimulation of the SSFO evoked neuronal activity, we performed both immunostaining and immunoblot analysis for c-Fos on brain sections or in hippocampal lysates, respectively. Coronal sections of mice infected with AAV-CaMKIIa-SSFO-mCherry in the hippocampus showed mCherry labelling in the CA1, CA2 and CA3 (Fig. 1B). Thirty minutes after light stimulation of mice expressing the SSFO-mCherry, the ipsilateral CA1, CA3 and dentate gyrus (DG) neurons displayed increased c-Fos as compared to neurons of the contralateral side (Fig. 1C). In mice infected with AAV-CaMKIIa-tdTomato, neurons in both hippocampus sides were negative for c-Fos (Supplementary Fig. S1). The c-Fos expression levels were significantly higher in hippocampal lysates from mice expressing SSFO as compared to tdTomato expressing mice (Supplementary Fig. S2).

**Fig. 1:**
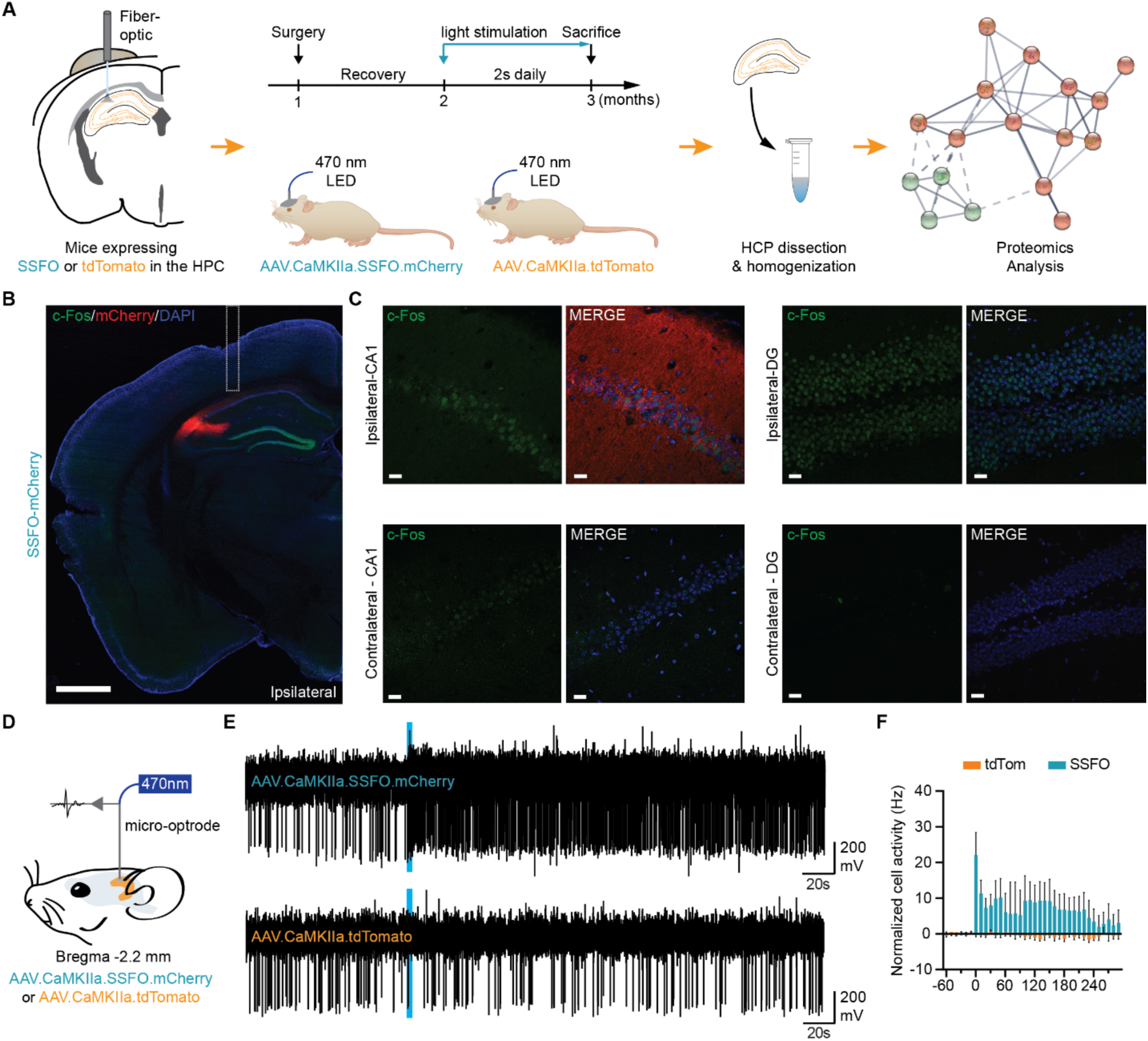
SSFO optogenetic stimulation activates hippocampal neurons and evokes neuronal hyperactivity. (**A**) Diagram showing an overview of the proteome experiment following chronic optogenetic stimulation of the hippocampus (HPC). (**B** - **C**) Confocal images showing immunofluorescence labeling of c-Fos (green) and mCherry (virus expression reporter protein, red) in coronal sections of mice infected with AAV-SSFO (Scale bars: 1 mm in **B** and 20 μm in **C**; DAPI in blue). Expression of SSFO was observed in the ipsilateral side of infection, but not the contralateral side. Unilateral light stimulation increased the levels of c-Fos at ipsilateral CA1 and DG but not the contralateral side. (**D**) Micro-optrode recording configuration. A 470 nm laser was coupled to an optrode probe for electrical recordings and advanced into the brain. (**E**) Example traces recorded in AAV-SSFO (top) or AAV-tdTomato (bottom) infected mice. 470 nm activation pulse is indicated with a blue bar. (**F**) Peristimulus time histogram of the mean cell activity of neurons recorded from mice infected with AAV-SSFO (n = 10; cyan) or AAV-tdTomato (n = 6; orange). Data presented as mean ± SEM.

Using extracellular recordings in anesthetized mice we assessed the light-induced changes in the activity of the CA1 region transduced either with the SSFO or tdTomato using a glass micro-optrode we previously designed, which enables dual, co-registered electrophysiological recording and optogenetic stimulation, free of photo-electric artefacts (Fig. 1D) (*25*, *26*). In mice expressing the SSFO, the multiunit spike rates within the CA1 increased for several minutes after a 2s light pulse as expected (Fig. 1E, F). Conversely, in tdTomato expressing mice, multiunit spiking activity did not increase following the brief light pulse (Fig. 1E, F). Altogether, these results indicate that the SSFO is transduced in the hippocampus and can be activated by light stimulation to generate neuronal hyperactivity.

### Chronic optogenetic stimulation of the hippocampus activates protein translation and phosphorylation but downregulates synaptic proteins in wild-type mice

We sought to identify and quantify alterations in protein levels in response to sustain neuronal hyperactivity in the hippocampus of mice. We analyzed the hippocampi of WT mice expressing SSFO or tdTomato after 4 weeks of daily brief light stimulation (Fig. 1A) using label-free liquid chromatography with tandem mass spectrometry (LC-MS/MS) proteomic analysis. We identified 778 proteins differentially abundant (P value ≤ 0.05) between the two conditions (Fig. 2A; Supplementary Table S1). The chronic optogenetic stimulation in mice expressing SSFO significantly upregulated the expression of 556 proteins and only downregulated the expression of 222 proteins (Fig. 2A). Gene Ontology (GO) enrichment analysis with GeneCodis (*27*) revealed that the abundant proteins were mainly part of the cytoplasmic and mitochondrial cellular components (Fig. 2B). Yet, 77 of the proteins identified were part of the synapse (Fig. 2B) with approximately 45% appearing downregulated, suggesting that synaptic function is altered by evoked neuronal hyperactivity.

**Fig. 2:**
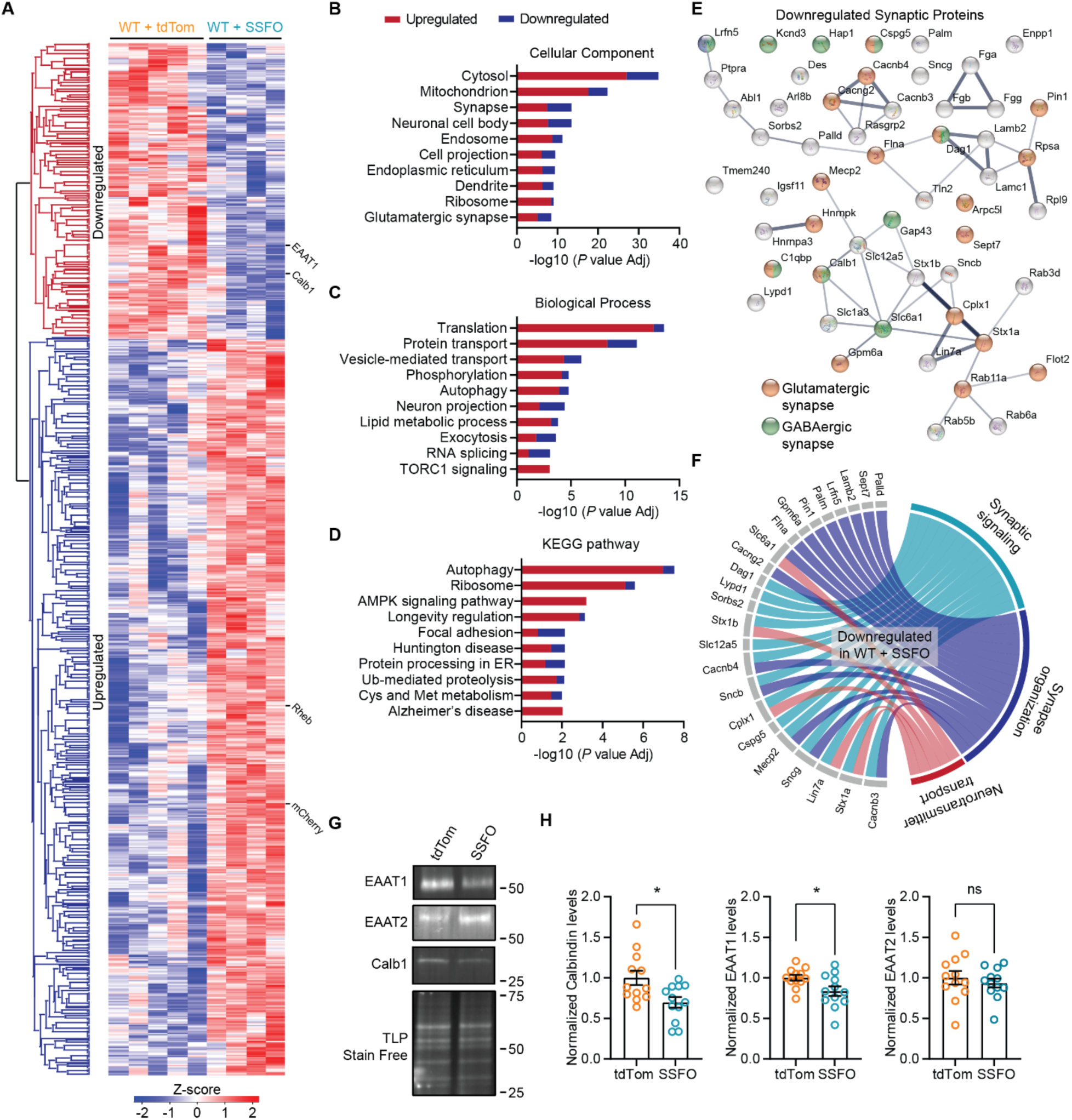
Chronic optogenetic stimulation elicits synaptic signalling protein downregulation in wild-type mice. (**A**) Heat map illustrating the clustering of proteins (*P* value ≤ 0.05) in the hippocampus of WT mice expressing SSFO or tdTomato after 4 weeks of daily optogenetic stimulation. The color-coded scale (Z-score) denotes the upregulated (red) and downregulated (blue) proteins. (**B** - **D**) Results of the GO enrichment analysis of the top 10 most abundant cellular components (**B**), biological processes (**C**) and KEGG pathways (**D**) in the hippocampus of WT+SSFO mice. In red or blue the fraction of the total proteins upregulated or downregulated, respectively, for each GO-term. (**E**) Interactome of functional or physical protein to protein associations of the synaptic proteins found downregulated in WT+SSFO mice. The proteins in orange are associated with glutamatergic synapses while the proteins in green with GABAergic. (**F**) Circos plots showing the proteins and selected biological processes downregulated in WT+SSFO mice. (**G**) Representative immunoblots for the levels of Calbindin (Calb1), EAAT1 and EAAT2 in hippocampal lysates from WT+SSFO and WT+tdTom mice. (**H**) Quantification of Calbindin, EAAT1 and EAAT2 protein levels measured by immunoblotting in WT+SSFO (cyan; *n* = 12) and WT+tdTom (orange; *n* = 12) mice after chronic optogenetic stimulation. The total protein (TLP) levels measured on TGX Stain-Free gels is the loading control.

Moreover, GO analysis revealed that translation, protein transport, phosphorylation and autophagy were among the most enriched biological processes found altered by the chronic optogenetic stimulation (Fig. 2C). In turn, the AMPK signaling, and ubiquitin-mediated proteolysis were identified among the top 10 most enriched KEGG pathways (Fig. 2D). Surprisingly, the AD pathway was also identified, with 31 proteins, all found upregulated in the WT+SSFO mice (Fig. 2D).

Out of the 34 downregulated synaptic proteins, interaction network analysis with STRING (*28*), identified 19 proteins as glutamatergic synapse proteins, while 9 were enriched in GABAergic synapses (Fig. 2E). And yet there are several more proteins, key for synaptic transmission, which were found significantly downregulated in WT mice following chronic hippocampal hyperactivity (Fig. 2E, F). Using immunoblots, we confirmed that Calbindin 1 (Calb1) was indeed downregulated in hippocampal lysates from WT+SSFO expressing mice as compared to WT+tdTom mice (Fig. 2G, H). Additionally, the excitatory amino acid transporter 1 (EAAT1 or Slc1a3) was downregulated, while the excitatory amino acid transporter 2 (EAAT2 or Slc1a2) was unaltered in the WT+SSFO mice (Fig. 2G, H), consistent with the findings from the proteomic analysis (Fig. 2A; Supplemental Table S1).

### Early AD pathology in 5xFAD mouse hippocampus largely mimics the footprint of altered proteostasis caused by chronic optogenetic stimulation

The 5xFAD mice exhibit amyloid deposition accompanied by cognitive deficits recapitulating major features of Alzheimer’s disease amyloid pathology (*29*). Comparison of AD human and rodent brain tissue revealed that the 5xFAD mice exhibit a proteomic signature similar to symptomatic AD (*30*). Moreover, an independent analysis of the mouse hippocampal proteome focused on the proteomic changes during progression of the amyloid pathology (*31*). In turn, we analyzed the proteome changes manifesting in 3-month-old 5xFAD mice, to identify similarities between the pre-symptomatic AD brain and the effect of hippocampal neuronal hyperactivity in WT mice.

When we compared the proteins altered by the mutations and the AD pathology in the hippocampus of 5xFAD mice, we identified 926 proteins as differentially expressed (Fig. 3A; Supplementary Table S2). The majority of these proteins were upregulated (Fig. 3B). Protein translation and transport, autophagy and phosphorylation were identified among the altered biological processes in the 5xFAD mice (Fig. 3C), consistent with our findings in the WT+SSFO mice (Fig. 2C). The mitochondria, synapse and endosomes were among the most enriched cellular components (Fig. 3D). The majority of the abundantly expressed synaptic proteins belong to the glutamatergic synapse, with 61% of them found downregulated in 5xFAD mice (Fig. 3D).

**Fig. 3:**
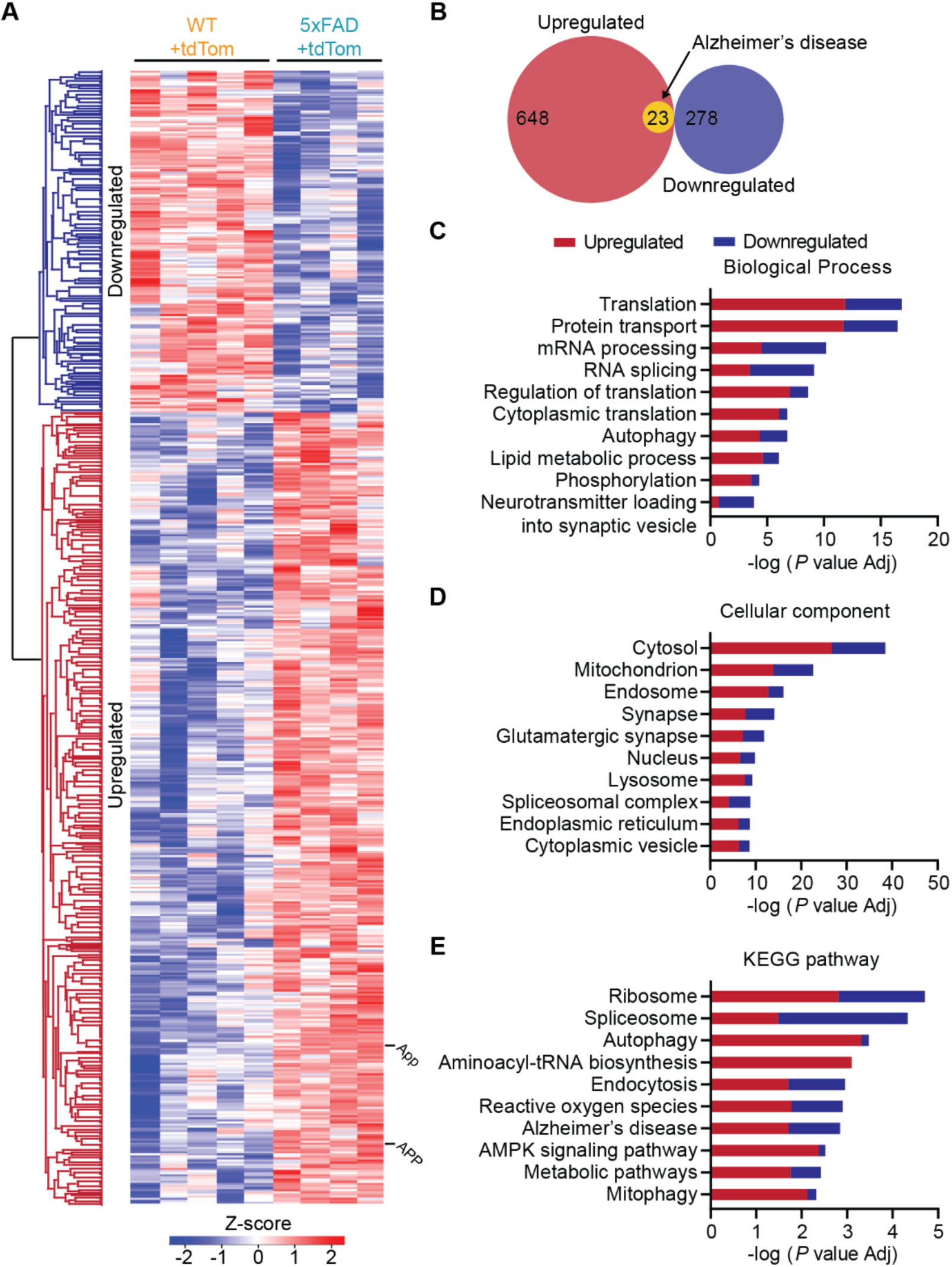
Early AD pathology alters translation, phosphorylation and autophagy in the hippocampus of 5xFAD mice. (**A**) Heat map illustrating the scaled protein levels in the hippocampus of 3-month-old 5xFAD mice. The color-coded scale (Z-score) denotes the upregulated (red) and downregulated (blue) proteins. (**B**) Venn diagram showing the number of upregulated (red) and downregulated (blue) proteins in 5xFAD mice. In yellow the number of upregulated proteins found enriched in the AD KEGG pathway. (**C - E**) The results of the GO enrichment analysis of the top 10 most abundant biological processes (**C**), cellular components (**D**) and KEGG pathways (**E**) in the hippocampus of 5xFAD mice. The colors in each bar represent the fraction of the total proteins for each GO-term found upregulated (red) or downregulated (blue).

Interestingly, for biological processes such as translation, phosphorylation, autophagy and RNA slicing the common proteins between the WT+SSFO and 5xFAD mice showed more than 50% overlap between the two groups (Fig. 4A), as well as the same valence (Fig. 4B). For the proteins found in the synapse the overlap was at 45.5% (Fig. 4A) and only Arr3 showed contrasting valence between the two groups (Fig. 4B). Finally, the GO analysis identified 38 proteins implicated in the AD pathway in the presymptomatic 5xFAD mice (Fig. 3E, 4D), with 19 of them also found abundantly expressed in WT+SSFO mice (Fig. 4B, C, E), indicating that chronic optogenetic activation of the hippocampus in WT mice induced signalling cascades involved in the AD pathogenesis.

**Fig. 4:**
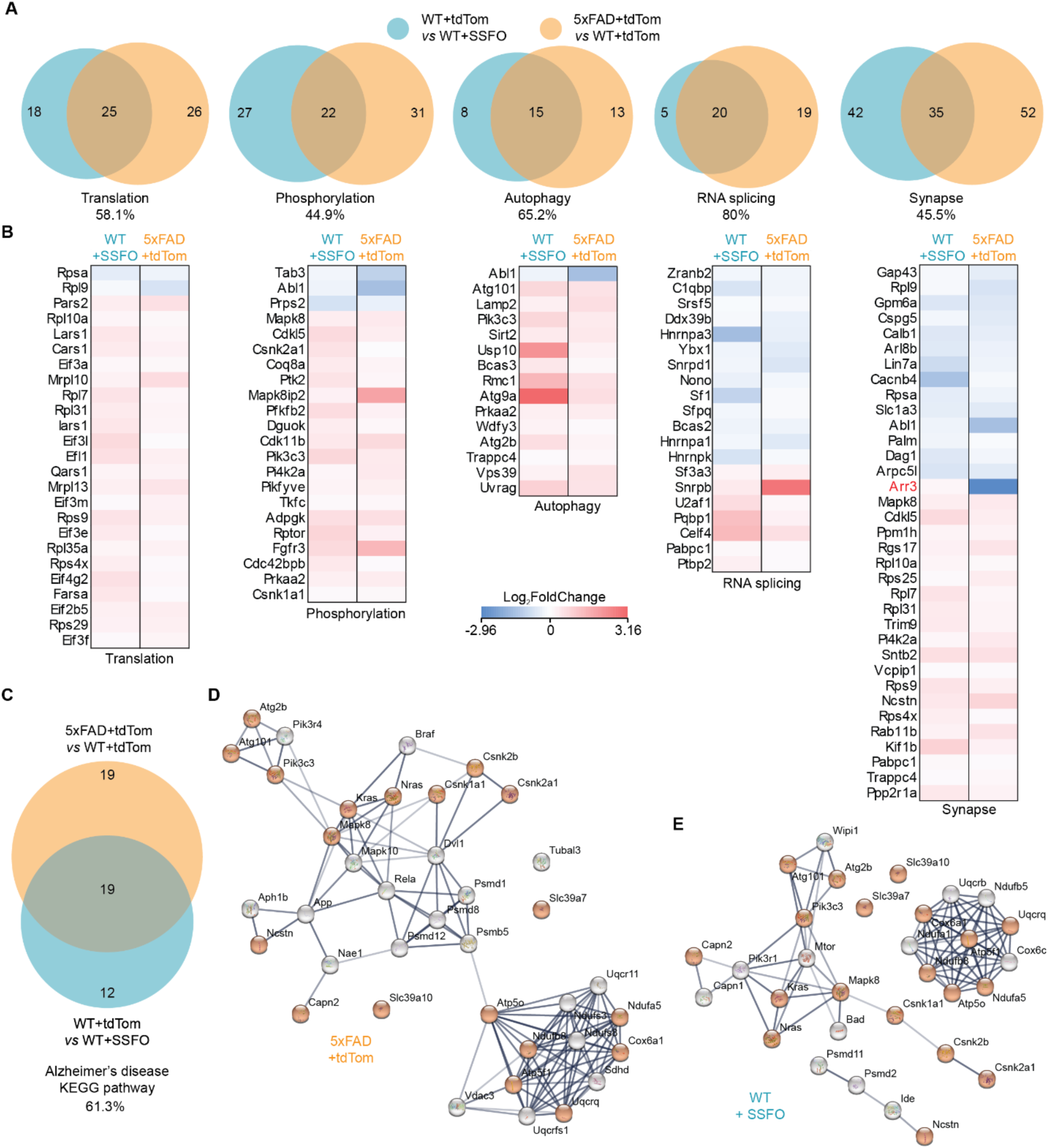
Chronic optogenetic activation of the hippocampus in wild-type mice induces proteome changes involved in the early AD pathogenesis. (**A**) Venn diagrams illustrating the number of the abundant proteins for each GO-term found in the hippocampus of WT+SSFO and 5xFAD+tdTom mice. The percentages represent the fraction of the WT+SSFO altered proteins commonly found altered in the 5xFAD mice. (**B**) Heat maps revealing the fold of change in levels for the common proteins of each GO-term in **A** in the WT+SSFO or 5xFAD+tdTom mice. (**C**) The number of common proteins participating in the AD KEGG pathway when comparing the altered proteins in the 5xFAD mice and the proteins altered in the WT+SSFO after the chronic optogenetic stimulation. (**D - E**) Interactomes of functional or physical protein to protein associations of the AD KEGG pathway proteins found altered in 5xFAD+tdTom (**D**) and WT+SSFO (**E**) mice.

### Evoked chronic neuronal hyperactivity in the hippocampus of 5xFAD mice downregulates mRNA splicing and protein phosphorylation

We adopted the same approach to induce neuronal hyperactivity in the hippocampus of 5xFAD mice (Fig. 1A). The chronic optogenetic stimulation in 5xFAD mice resulted in significantly downregulating the expression levels of 264 proteins while it resulted in upregulating the levels of only 99 proteins (Fig. 5A; Supplementary Table S3). The altered proteins were primarily cytoplasmic, nucleic and part of the spliceosomal complex (Fig. 5B). Among the cellular components greatly enriched, the glutamatergic synapse was identified with 29 proteins (Fig. 5B), 83% of which were downregulated. Three additional proteins of the GABAergic synapse were found downregulated in the 5xFAD mice after the chronic optogenetic stimulation (Fig. 5C).

**Fig. 5:**
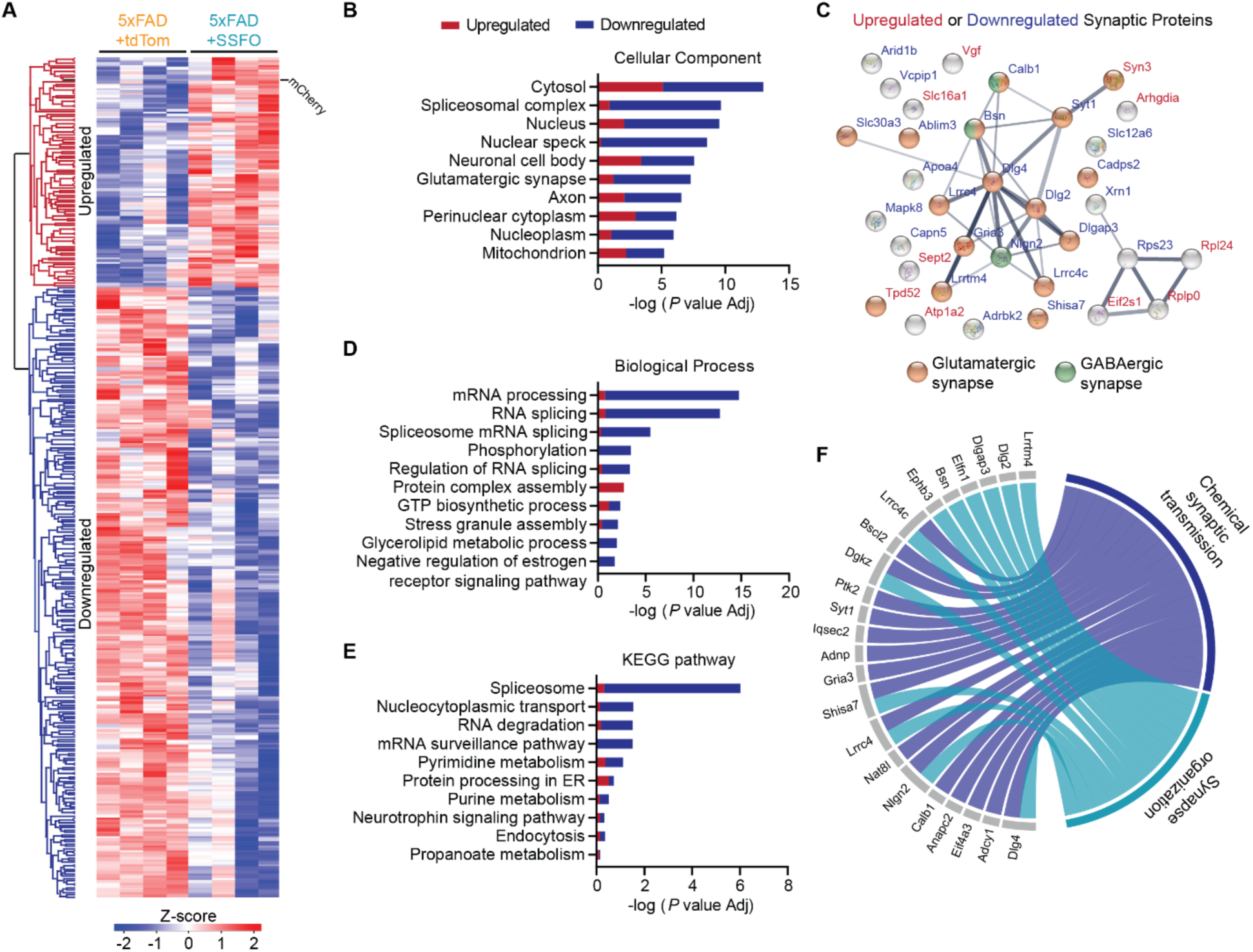
Chronic optogenetic stimulation downregulates RNA processing and protein phosphorylation in 5xFAD mice. (**A**) Heat map revealing the scaled protein levels in the hippocampus of 5xFAD mice expressing SSFO or tdTomato after 4 weeks of daily optogenetic stimulation. (**B**) GO enrichment analysis showing the top 10 most abundant biological processes. In red or blue the fraction of the total proteins upregulated or downregulated for each GO-term, respectively (**C**) Interactome of functional or physical protein to protein associations of the synaptic proteins altered in 5xFAD+SSFO mice after optogenetic stimulation. The orange circles represent glutamatergic synapse proteins while the green GABAergic. The proteins written in red are upregulated while the proteins in blue are downregulated. (**D** - **E**) Same as in **B** but for the top 10 cellular components (**D**) and KEGG pathways (**E**) in the hippocampus of 5xFAD+SSFO mice. (**F**) Circos plots showing selected biological processes and the proteins altered in 5xFAD+SSFO mice.

Contrary to the WT mice, GO enrichment analysis revealed that neurons in 5xFAD mice responded distinctly to the induced neuronal hyperactivity by downregulating mRNA processing, splicing and protein phosphorylation (Fig. 5D). Consistently, the spliceosome and RNA degradation appeared among the most enriched KEGG pathways in the 5xFAD+SSFO mice (Fig. 5E). The purine and pyrimidine metabolism pathways were also affected (Fig. 5E). Lastly, biological processes such as chemical synaptic transmission and synapse organization appeared altered (Fig. 5F) by the chronic optogenetic stimulation in the 5xFAD mice. These findings indicate that the 5xFAD mice respond distinctively to the chronic stimulation, as compared to the WT mice, and thus the plasticity mechanisms induced by the stimulation in WT mice matches the changes occurring at early rather than late stages of the AD pathogenesis.

### Degree of similarity in proteome changes caused by hyperactivity in WT brain *vs*. that in AD-related pathology

Hippocampal hyperactivity has been implicated in memory deterioration and AD pathogenesis (*5*, *12*). To further explore whether chronic optogenetic activation in the hippocampus of WT mice indeed replicates some of the neurodegenerative processes involved in the pathogenesis of AD, we performed a hierarchical clustering analysis based on our proteomic data. Accordingly, we found that the WT+SSFO mice grouped together with the 5xFAD+tdTom while all the WT+tdTom mice separated from the rest of mice according to the changes in protein levels (Fig. 6A). Moreover, when we compared the proteins altered by stimulation in WT+SSFO mice to the ones altered due to the mutations and AD-related pathology in the 5xFAD mice, we found that those two groups resemble themselves much more than when comparing 5xFAD mice to unstimulated control (only 135 protein changes in common; Fig. 6B).

**Fig. 6:**
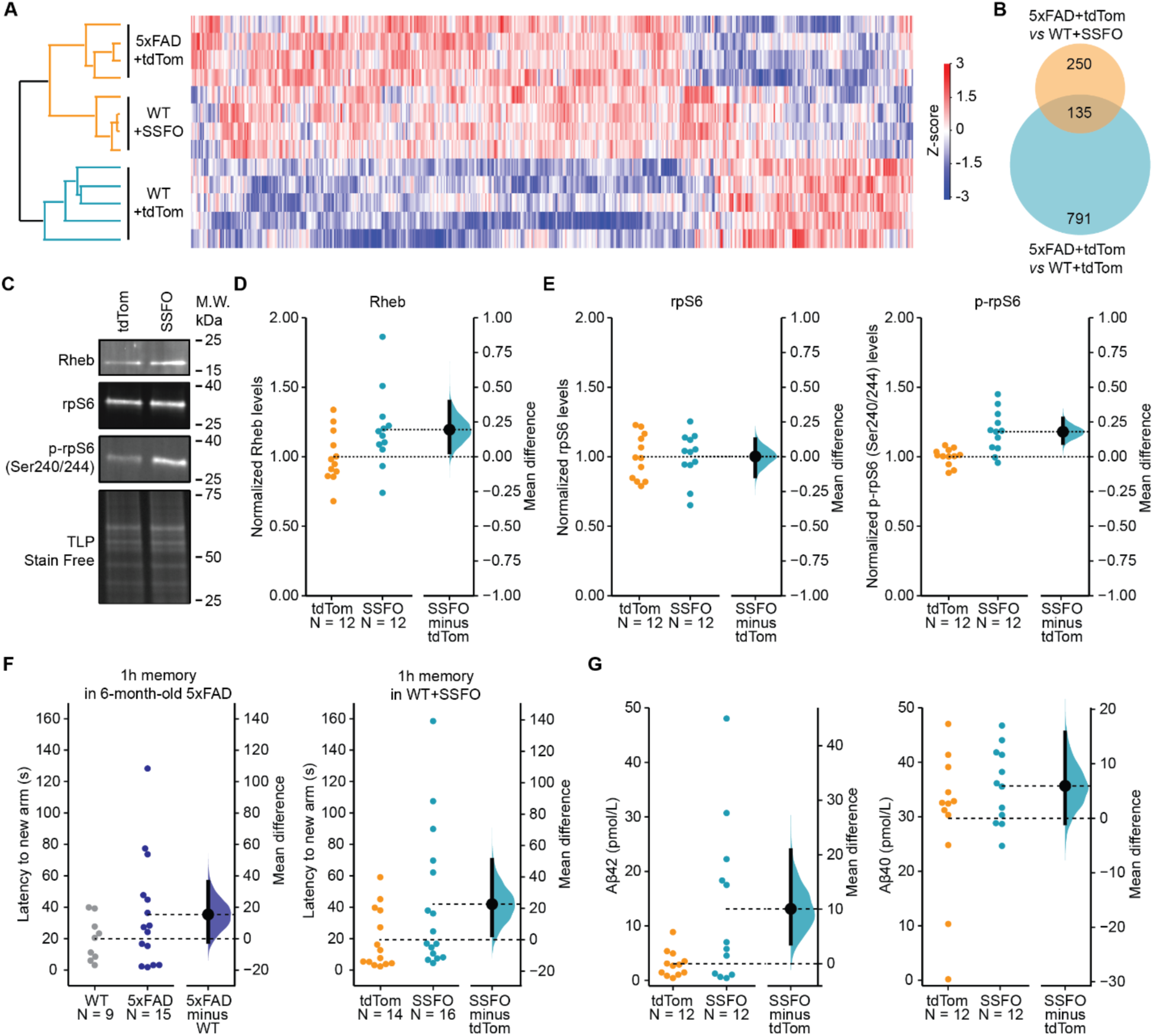
Chronic hippocampal optogenetic activation induces spatial memory deficits and augments Αβ42 secretion in wild-type mice. (**A**) Heat map revealing the hierarchical clustering of 5xFAD and WT mice expressing SSFO or tdTomato based on their scaled protein levels in the hippocampus after 4 weeks of daily optogenetic stimulation. (**B**) Venn diagram showing the number of common proteins when comparing the altered proteins in 5xFAD mice as compared to WT (5xFAD+tdTom *vs* WT+tdTom) and the proteins altered between the 5xFAD+tdTom and the WT+SSFO. (**C**) Representative immunoblots of hippocampal lysates from WT+SSFO and WT+tdTom mice. (**D**) The protein levels of Rheb in WT+SSFO and WT+tdTom mice. (**E**) Quantification of ribosomal protein S6 (rpS6) expression (left) and its phosphorylation levels at the Ser240/204 residues. (**F**) The latency to the new arm during the one-hour Y-maze spatial memory test for 6-month-old 5xFAD mice (*n* = 15) *vs* their non-transgenic littermates (WT; *n* = 9) on the left and for WT+SSFO mice (*n* =12) *vs* WT+tdTom mice (*n* = 12) on the right. (**G**) The concentration of soluble Αβ42 (left) and Αβ40 (right) in the hippocampus of WT+SSFO mice as compared to WT+tdTom. The data are analyzed with estimation statistics. Each point represents an individual mouse. The mean difference is dissipated as a dot and the 95% confidence interval by the ends of the vertical error bars.

The mammalian target of rapamycin complex 1 (mTORC1) is a principal signalling pathway controlling protein translation initiation (*32*) and has been postulated to take part in AD pathology (*33*). Immunoblotting of hippocampal lysates revealed an increase in the phosphorylation of the mTORC1 downstream translation initiation effector ribosomal protein S6 (rpS6; Fig. 6C, E), concomitant with an increase in the expression of Rheb (Fig. 6C, D), the mTORC1 canonical activator (*34*) in the WT+SSFO mice as compared to the WT+tdTom mice.

### Neuronal hyperactivity in wild-type mice induces spatial memory deficits and augments Αβ42 secretion

To further examine how chronic hippocampal hyperactivity and the synaptic transmission changes (Fig. 2F) observed following the optogenetic stimulation impact brain function, we performed behavioral testing of WT and 5xFAD mice. We adopted a modified version of the Y-maze alternation test allowing for the evaluation of spatial memory performance in mice (Supplementary Fig. S3). We found that chronic optogenetic stimulation in the hippocampus of WT+SSFO mice resulted in spatial memory deficits, similar to the impairment seen here in 6-months-old 5xFAD mice (Fig. 6F), or previously described in (*35*).

Finally, since previous observations have linked neuronal and network hyperactivity with an increase in the secretion and deposition of amyloid-β in the brain of transgenic mice carrying several AD-linked mutations (*17*, *18*, *36*, *37*), we sought to measure the levels of soluble Αβ40 and Aβ42 in the brain of WT mice following our chronic optogenetic stimulation protocol. In the hippocampus of WT+SSFO mice the levels of soluble secreted Aβ42 were significantly increased as compared to the levels in the brain of WT+tdTom mice (Fig. 6G). The levels of Aβ40 showed a trend of increase in the WT+SSFO mice, but this increase was not significantly different to the Αβ40 levels in the WT+tdTom (Fig. 6G). Altogether, these results suggest that chronic neuronal hyperactivity is sufficient to induce synaptic transmission disruption and memory loss similar to that observed in the neurodegenerative process in AD, but also augments the amyloidogenic cleavage of App.

## Discussion

We adopted optogenetics and an SSFO to induce chronic neuronal hyperactivity and examine the effect of such manipulation in the hippocampus of young WT and 5xFAD mice. We present evidence of (i) an overt downregulation in the levels of both excitatory and inhibitory synaptic proteins and an activation of translation, phosphorylation and autophagy following neuronal hyperactivity in WT mice; (ii) an upregulation of AD-associated proteins in WT+SSFO mice mimicking the AD proteome signature found in the 5xFAD mice; (iii) a downregulation in the levels of proteins participating in RNA processing and protein phosphorylation in 5xFAD+SSFO mice; (iv) spatial memory impairments in WT+SSFO mice upon light stimulation; and (5) elevated levels of soluble Αβ42 in WT hippocampi following chronic optogenetic stimulation. These findings strongly support the notion that rampant neuronal hyperactivity distorts cellular mechanisms regulating proteostasis similar to that seen in the prodromal phase of AD-like pathology. This positions overt neuronal hyperactivity as a substrate of AD pathogenesis.

In various neurological disorders, cognitive decline has been associated with hippocampal hyperactivity and the failure to deactivate brain regions consisting of the default mode network (e.g. posterior cingulate cortex, precuneus, retrosplenial cortex) (*5*, *38–42*). Reducing neuronal hyperactivity pharmacologically has been also found to improve cognitive performance in humans with mild cognitive decline (*43*, *44*) suggesting that neuronal hyperactivity is a cause and not a compensatory mechanism of cognitive decline. But neuronal hyperactivity may have broader implications in disease onset. Apart from contributing to cognitive deficits, neuronal hyperactivity could drive several other pathological aspects related to AD, such as mTORC1 hyperactivity (*45*), Αβ release (*18*), synaptic loss (*46*, *47*) and neuronal degeneration (*38*).

It has been shown before that electrical, pharmacological and optogenetic neuronal stimulation can augment the secretion of Αβ from the presynaptic terminals in various APP transgenic mouse lines (*17*, *18*, *36*, *37*). In turn, elevated Αβ levels have been shown to induce neural overexcitability and hyperactivity disrupting neural circuits both in WT and APP transgenic mouse lines (*10*, *13*, *48*). Moreover, in humans, within the brain regions where Αβ aggregation is pronounced, the basal metabolic rates and neuronal activity are profoundly elevated (*8*, *37*, *49*). These findings support the hypothesis of a vicious unclamped cycle facilitating an Αβ-dependent neuronal hyperactivity in AD (*13*) and are in line with our results showing an increase in soluble Αβ42 secretion and a multifactorial perturbation of synaptic transmission in WT animals following neuronal hyperactivity.

The similarity we found in altered proteostasis and spatial memory in overstimulated WT and 5xFAD mice, together with the finding that neuronal hyperactivity in WT mice augments Αβ42 secretion establish chronic optogenetic stimulation in WT animals as a potential paradigm to model sporadic AD-pathogenesis.

Finally, the distinct response to the chronic optogenetic stimulation in the 5xFAD, at least at the proteome level, may reflect an occlusion of the detrimental effects of evoked neuronal hyperactivity in AD, or a loss of proteostasis reserve. This is in line with observations indicating that neuronal hyperactivity manifests early in AD resulting in synapse and circuit disruption (*5*, *6*, *50*, *51*). And yet, while these deficits in circuits arising from neuronal degeneration in AD might appear beyond repair, suppressing excessive neuronal activity early in the progression of the disease may represent a possible therapeutic modality to delay or modify the disease progression and cease the vicious cycle of relentless hyperactivity.

## Materials and Methods

### Animals

Rodent experiments were carried out on male 5xFAD mice, and their WT littermates bred and housed in the CERVO Brain Research Centre animal facility. The male 5xFAD and female B6SJLF1/J breeders were purchased from the Jackson Laboratory (#34840-JAX and #100012, respectively). Mice were housed on a 12 h day/night cycle with ad libitum access to food and water. Mice were randomly assigned to experimental groups. All experiments were approved by the committee for animal protection of Université Laval (CPAUL) and followed the guidelines from the Canadian Council for Animal Care.

### Surgical Procedures and viral transduction

All viral constructs were produced by the CERVO Canadian Optogenetics and Vectorology Foundry Core Facility (RRID: SCR_016477). The tip of the optic fiber (200 μm core, 0.39 NA on 2.5 mm ceramic ferrules) was coated with 100 nL of a 1:1 mix of silk fibroin solution (Sigma 5154-20ML) and the viral vector AAV2/5-CaMKIIa-hChR2(C128S/D156A)-mCherry (henceforth referred to as AAV2/5-CaMKIIa-SSFO-mCherry; titer: 7.7 x 10^12^ GC/mL) or AAV2/8-CaMKIIa-tdTomato (titer: 1.3 x 10^13^ GC/mL), prior to the surgery, and left to dry for 15 hrs at 4 °C as described in (*52*).

Stereotaxic surgeries were performed on mice (4-weeks old) maintained on isoflurane anaesthesia (1.5%-2% in O2). Briefly, we removed the skin atop the cranium and a ∼250-μm-diameter craniotomy was drilled over the hippocampus at coordinates relative to Bregma of −2.30 mm rostro-caudal and −2.00 mm latero-medial. The optic fiber cannula was lowered 1.60 mm into the brain (−1.60 mm dorso-ventral) and fixed on the cranium with C&B Metabond® (Parkell Inc.).

Following the surgery, mice were single-housed, and the chronic optogenetic stimulation experiments started 4-5 weeks later. For the optrode recordings, the same viral vectors were unilaterally infused into each side of the hippocampal CA1 (rostro-caudal: −2.30 mm, latero-medial: 2.00 mm, dorso-ventral: 1.60 mm). The infusion was performed with pulled borosilicate glass capillaries and a NANOLITER2020 injector (World Precision Instruments LLC). The in vivo optrode experiments started a minimum of 4 weeks later.

### Optogenetic stimulation

After 4-5 weeks from the optic fiber implantation mice were optically stimulated for 2 s every 24 hrs (starting at 2-months of age) for 4 weeks with blue light (470 nm, 1 mW). The light was generated by an LED diode (ThorLabs M470F3) and delivered through fiber optics (200 μm core, 0.39 NA) to the implanted fiber optic cannulae.

### *In vivo* extracellular optrode recordings

Simultaneous optical stimulation and electrical extracellular recording in the hippocampal CA1 was performed in WT male mice previously transduced with SSFO or tdTomato using a micro-optrode (*25*). Briefly, mice were anesthetized with a mix of 100 mg ketamine, 15 mg xylazine and 2.5 acepromazine per kg and placed in a stereotaxic frame on a temperature control pad. A craniotomy was drilled over the hippocampus at coordinates relative to Bregma of −2.30 mm rostro-caudal and −2.00 mm latero-medial. A micro-optrode (6-8 μm tip diameter; 550 μm optical core; 0.29 NA; 250 μm hollow core) was filled with 0.5 M potassium acetate solution, connected to a 488 nm laser diode (Doric lenses) through a multimode optical fiber (200 μm optical core; 0.22 NA; ThorLabs) and lowered vertically to the brain. Electrophysiological recordings were initiated at −1,3 mm ventral. A 2s long light pulse was delivered to activate the SSFO. The extracellular electrophysiological signal was amplified (Neurodata IR183, Cygnus technology), filtered (band pass, 300–3,000 Hz, model 440, Brownlee Precision), and digitized. The filtered traces were analyzed with Spike2 (Cambridge Electronic Design). Events were detected and the firing rate of individual neurons was plotted in peristimulus time histograms (PSTH) with 10 s bins. Neurons for which the standard deviation of firing rate for the light pulse bin was 1.5 times superior to the basal firing rate (average of the firing rate during the 60s preceding the pulse of light) were considered activated.

### Sample Preparation

For the proteomics analysis, brains were excised from anesthetized mice (30% urethane in saline) within 30 min after optogenetic stimulation. The hippocampi were dissected, collected in sample tubes, frozen immediately in liquid nitrogen, and stored at −80 °C until use. 25 mg of frozen tissue was lysed in 150 μL of freshly made lysis buffer (4% SDS, 0.1 M DTT, 100 mM Tris/HCl; pH 7.6) and homogenized with an electric FisherBrandTM Pellet PestleTM homogenizer (Thermo Fisher Scientific). Homogenates were incubated at 95 °C for 3 min and centrifuged at 16,000 g for 5 min. The supernatants were transferred into sample tubes and stored at −80 °C.

For ELISA and immunoblots, tissue was harvested as before within 30 min after optogenetic stimulation. 25 mg of frozen hippocampus tissue was homogenized in 150 μL of RIPA buffer (50 mM Tris-HCl, 150 mM NaCl, 1% Triton X-100, 0.5% Na deoxycholate, 0.1% SDS; pH 8.0). Homogenates were incubated under constant agitation for 4 hrs at 4 °C for lysis. Lysates were centrifuged at 17,200 g and 4 °C for 10 min. Supernatants were collected and stored at −30 °C.

### Protein mass spectrometry

Protein mass spectrometry analysis was performed in the Proteomics Facility of Biomedical Sciences Research Centre Alexander Fleming. Four to five biological replicas for each condition (WT + AAV2/8-CaMKIIa-tdTomato; WT + AAV2/5-CaMKIIa-SSFO-mCherry; 5xFAD + AAV2/8-CaMKIIa-tdTomato; 5xFAD + AAV2/5-CaMKIIa-SSFO-mCherry) were analyzed. Briefly, proteins of the lysed samples were processed according to the sensitive Sp3 protocol (*53*). The reduced cysteine residues were alkylated in 200 mM iodoacetamide (Acros Organics). 20 μg of beads (1:1 mixture of hydrophilic and hydrophobic SeraMag carboxylate-modified beads; GE Life Sciences) were added to each sample in 50% ethanol. Protein clean-up was performed on a magnetic rack. The beads were washed twice with 80% ethanol followed by one wash with 100% acetonitrile (Fisher Chemical). The beads-captured proteins were digested overnight at 37 °C with 0.5 μg trypsin/LysC mix in 25 mM ammonium bicarbonate under vigorous shaking (1200 rpm, Eppendorf Thermomixer). The supernatants were collected, and the peptides were purified by a modified Sp3 clean-up protocol and finally solubilized in the mobile phase A (0.1% formic acid in water), and sonicated. Peptides concentration was determined through absorbance measurement at 280 nm using a nanodrop instrument.

Peptides were analyzed by a liquid chromatography tandem mass spectrometry (LS-MS/MS) on a setup consisting of a Dionex Ultimate 3000 nanoRSLC online with a Thermo Q Exactive HF-X Orbitrap mass spectrometer. Peptidic samples were directly injected and separated on a 25 cm-long analytical C18 column (PepSep, 1.9 μm^3^ beads, 75 μm ID) using a 90 min long run. The full MS was acquired in profile mode using a Q Exactive HF-X Hybrid Quadropole-Orbitrap mass spectrometer operating in the scan range of 375-1400 m/z using 120 K resolving power with an Automatic Gain Control (AGC) of 3 x 10^6^ and a max IT of 60 ms followed by data independent analysis (DIA) using 8 Th windows (39 loops counts) with 15 K resolving power with an AGC of 3 x 10^5^, a max IT of 22 ms and normalized collision energy (NCE) of 26.

### Label-free quantification and data analysis

Orbitrap raw data were analyzed in DIA-NN 1.8 (Data-Independent Acquisition by Neural Networks) against the complete Uniprot Mus musculus proteome (Downloaded April 16, 2021) supplemented with APP, presenilin, mCherry and ChR2. Search parameters were set to allow up to two possible trypsin/P enzyme missed cleavages. A spectra library was generated from the DIA runs and used to reanalyze them. Cysteine carbamidomethylation was set as a fixed modification while N-terminal acetylation and methionine oxidations were set as variable modifications. The match between runs (MBR) feature was used for all the analyses and the output (precursor) was filtered at 0.01 false discovery rate (FDR). The protein inference was performed on the gene level using only proteotypic peptides. The double pass mode of the neural network classifier was also activated. Perseus (version 1.6.15.0) (*54*) was used for data processing and statistical analysis. Proteins were subjected to filtering based on valid values setting a threshold of 70% valid values in at least one group. Remaining missing values were imputed separately for each column (based on the normal distribution using width of 0.3 and down shift of 1.8). The statistically significant proteins were then Z-scored and visualized by Euclidean hierarchical clustering as heat maps.

Gene-Ontology analysis was performed with GeneCodis (*27*) by uploading the list of altered proteins based on the T-test two-sample comparisons. The interaction network analysis of functional or physical protein to protein associations were performed with STRING (version 11.5) (*28*).

### Immunoblots

Hippocampal lysates were diluted in Laemmli buffer at a concentration of 10 μg per 20 μL and denaturized for 5 min at 95 °C. Proteins were separated in 4–15% Mini-PROTEAN® TGX Stain-Free™ acrylamide gels (BioRad) and transferred onto 0.45 μm low-fluorescence PVDF membranes (BioRad). Membranes were probed with the rabbit monoclonal anti-EAAT1 (Abcam ab176557), anti-EAAT2 (Abcam ab205248), anti-S6 (Cell Signaling 2217), anti-S6 Ser240/244 (Cell Signaling 5364), anti-Rheb (Cell Signaling 13879), anti-c-Fos (Cell Signaling 2250) or the rabbit polyclonal anti-Calbindin (Millipore ABN2192) in Tris-buffered saline containing 0.2% Tween and 5% bovine serum albumin at 4 °C. All antibodies were used at 1:1000, except from the anti-S6 Ser240/244 which was used at 1:10000. The appropriate anti-mouse or anti-rabbit IgG conjugated with IRDye©800 or IRDye©680, respectively, were used at 1:10000. Proteins were visualized using a ChemiDoc Imaging System (BioRad) and the intensity of protein bands was quantified using ImageJ (NIH). To normalize for sample loading, the intensity of total protein (TLP) measured on the TGX Stain-Free™ gels was used as previously described (*55*).

### ELISA

Levels of Αβ40 and Aβ42 in hippocampal samples were quantitated by Human/Rat β Amyloid (40) ELISA kit (Wako 294-62501, LOT# WTL5240) and Human/Rat β Amyloid (42) ELISA kit (Wako 290-62601, LOT# WTM4353) respectively. Both ELISAs were performed according to the manufacturer recommendations, and the plates were read at 450 nm using an Eon microplate reader (BioTek).

### Immunofluorescence

A Leica Vibratome VT1220S (Leica Microsystems) was used to cut 100 μm coronal sections of paraformaldehyde fixed brain tissue. Sections were rinsed 3 times in 0.1M phosphate-buffered saline with 0.2% Triton X-100 (PBST) for 10 min, blocked for 1 hr with 10% Normal Goat Serum (NGS) in PBST and then incubated for 15 hrs at 4 °C in primary anti-c-Fos (rabbit monoclonal, 1:2000, Cell Signaling 2250) diluted in PBST containing 4% NGS. Sections were washed in PBST and subsequently incubated for 2 hrs at room temperature in AlexaFluor™ 488-conjugated goat anti-rabbit (1:500, Invitrogen A11008) diluted in PBST containing 4% NGS. Tissue was mounted on SuperFrost™ slides (Thermo Fisher Scientific 12-550-15) using a fluorescence mounting medium with DAPI (Abcam ab104139) and cover-slipped.

All confocal images were acquired with a Zeiss LSM710 confocal laser scanning microscope. Acquisitions were 12-bit images, 2048×2048 pixels with a pixel dwell time of 3.15 μs. A 40x Plan-Apochromat oil objective (1.4 NA) was used for magnification. Tile scans were acquired at 12-bits, 512×512 pixels (each tile) with a pixel dwell time of 0.64 μs and an x5 EC-Plan-Neofluar objective (0.16 NA). Laser power, photomultiplier tube (PMT) settings, filters, dichroic mirrors, scanning speed were kept constant for all acquisitions.

### Y-maze spatial memory test

The Y-maze apparatus consisted of 3 identical arms (each 36.2 x 8.25 cm) in a Y configuration (placed at 120° to each other) connected by a center polygonal area. The walls of the arms were transparent so the animals could identify two distinct visual cues placed outside the apparatus to facilitate spatial navigation. During the training phase, the mice were placed individually in the maze and left freely to explore two arms, for 10 minutes, while the third arm was blocked by a removable door. An hour later, the mice were returned to the maze and allowed to explore all arms. The latency to enter the new (third) arm was measured for each mouse and used to score the spatial memory performance of the mice. Between mice, the maze was cleaned with 70% isopropyl solution to minimize scent cues. Entry into the arm was defined as a mouse placing all four paws on the arm. For the mice receiving the daily optogenetic stimulation, the spatial memory test was performed after the end of the 4 weeks-long stimulation protocol. The day of the test, the mice receive a 2s light pulse before the beginning of the training phase.

### Statistical Analysis

For the proteomic experiments, the statistical analysis of the label-free quantification intensities was performed with Perseus (version 1.6.15) using a two sample t-test with a P-value less than 0.05 or an ANOVA for multiple sample tests with a P-value lower than of 0.05. For estimation statistics based on effect size and confidence intervals (*56*), the data were uploaded and analyzed on https://www.estimationstats.com/. The estimation plots show the mean difference between the groups and the 95% confidence interval by the ends of the vertical error bar. Bar graphs were generated with GraphPad Prism 9 (GraphPad software). Circos plots were drawn using RStudio by uploading the list of proteins enriched in our samples for each biological process selected by the gene-ontology analysis.

## Acknowledgments

The authors thank Camille Sugère and Lyane Méthot for assistance with the daily optogenetic stimulation protocol and Drs Efthimios MC Skoulakis and Nicolò Ilacqua for their invaluable comments on the proteomics data. The authors also thank Dominique Isabel for assistance with RStudio and data visualization.

## Funding

Weston Family Foundation Transformation Research grant TR192089 (YDK)

Canadian Institutes of Health Research (CIHR) grant FDN - 159906 (YDK)

Canada Research Chair program (YDK)

Université Laval Norampac Research grant on Alzheimer’s disease and Related Diseases (IK)

Université Laval Fondation De La Famille Lemaire research grant on Alzheimer’s disease and Related Diseases (IK)

## Author contributions

Conceptualization: IK, KP, YDK

Methodology: IK, MS, JA, KP, YDK

Investigation: IK, MS, RS, JA

Visualization: IK, KP, YDK

Supervision: KP, YDK

Writing—original draft: IK, KP, YDK

Writing—review & editing: IK, MS, KP, YDK

## Data and materials availability

The mass spectrometry proteomics data have been deposited to the ProteomeXchange Consortium via the PRIDE (*57*) partner repository with the dataset identifier PXD044437. The code used to generate the circos plot is available upon request. Any additional information required to reanalyze the data reported in this paper is available from the corresponding author upon request.

## Supplementary Materials

### Supplementary Figures

**Fig. S1.**
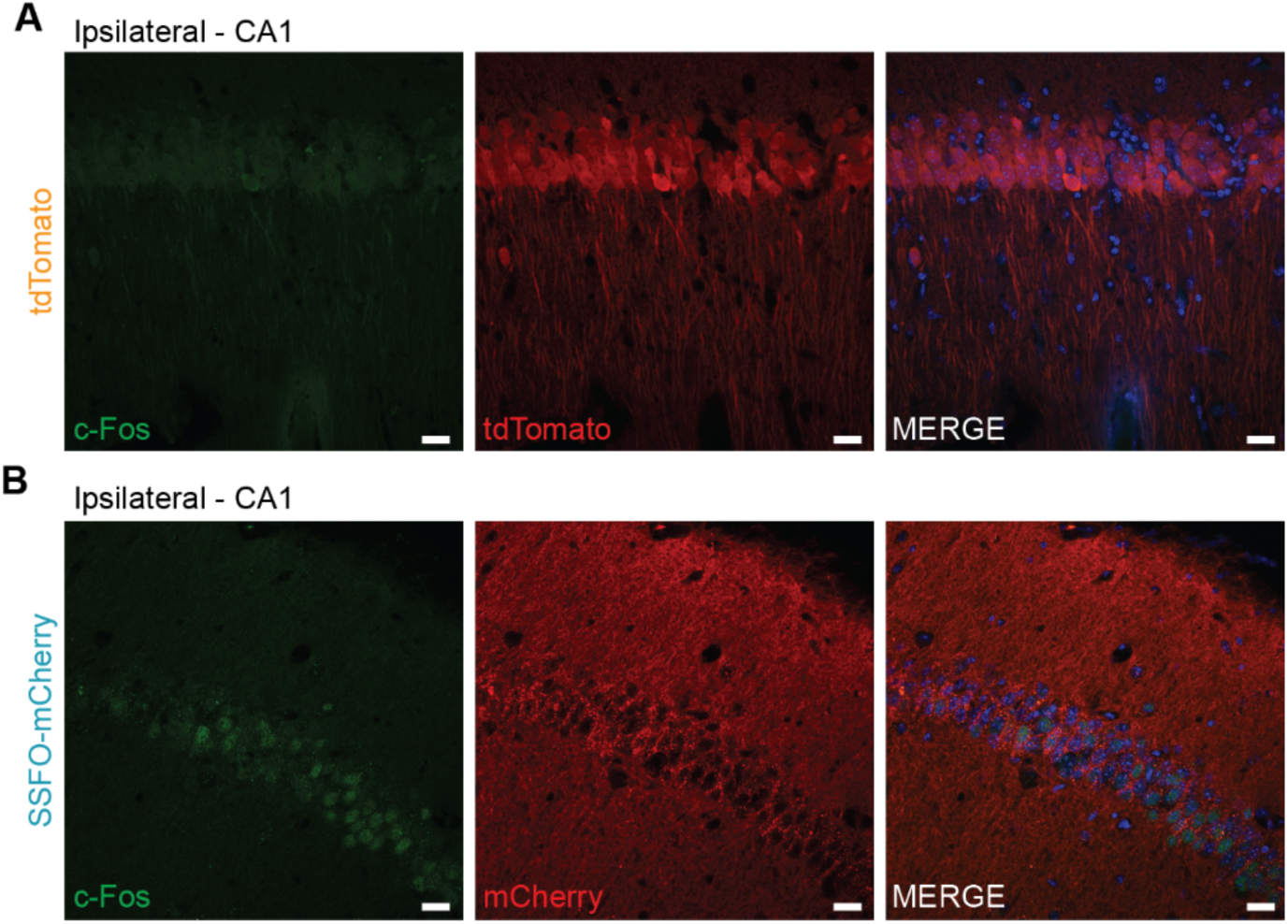
c-Fos expression in the hippocampus of AAV-CaMKIIa-tdTomato or AAV-CaMKIIa-SSFO-mCherry infected mice. (**A**) Confocal images showing labeling for c-Fos (green) and tdTomato (reporter protein, red) in coronal sections of mice infected with AAV-CaMKIIa-tdTomato. (**B**) Confocal images showing labeling for c-Fos (green) and Cherry (SSFO expression reporter protein, red) in coronal sections of mice infected with AAV-CaMKIIa-SFFO. Scale bars 20 μm, DAPI in blue.

**Fig. S2.**
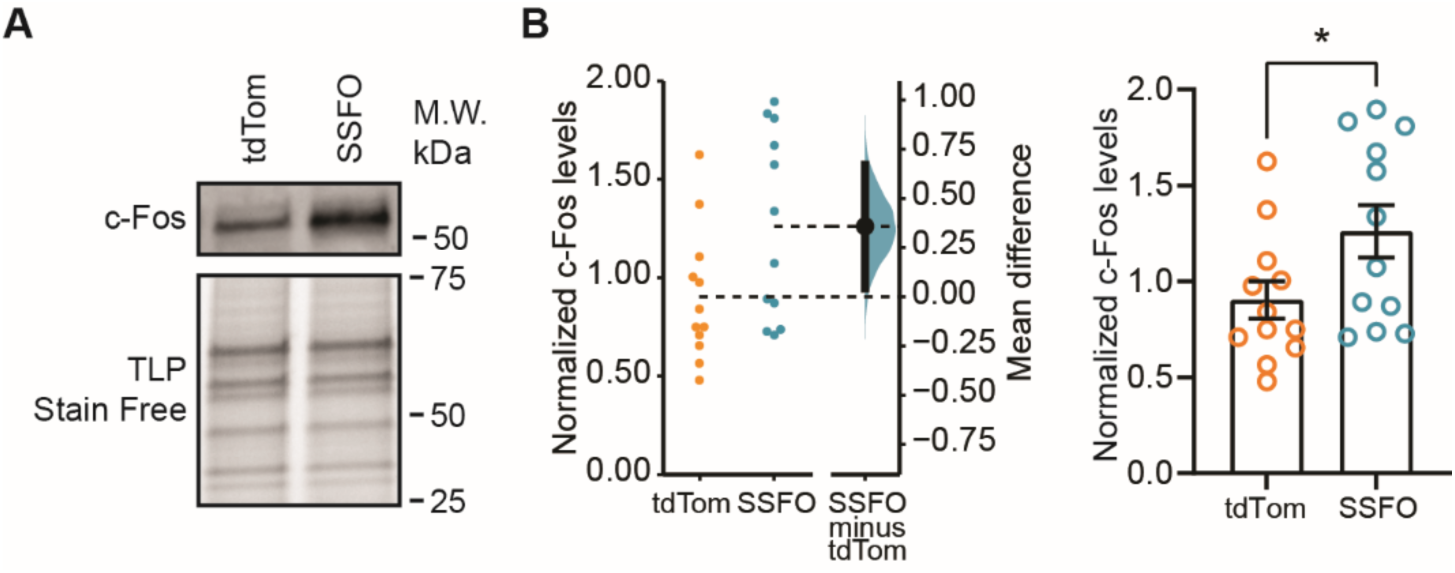
c-Fos is upregulated in the hippocampus of SSFO expressing wildtype mice. (**A**) Representative immunoblots for the levels of c-Fos in hippocampal samples from WT+tdTom and WT+SSFO mice. (**B**) Quantification of c-Fos protein levels measured by immunoblotting in WT+SSFO (cyan; n = 12) and WT+tdTom (orange; n = 12) mice. Data are represented as the mean difference (dot) and the 95% confidence interval (by the ends of the error bars; left) or as mean ± SEM (right). *P < 0.05.

**Fig. S3.**
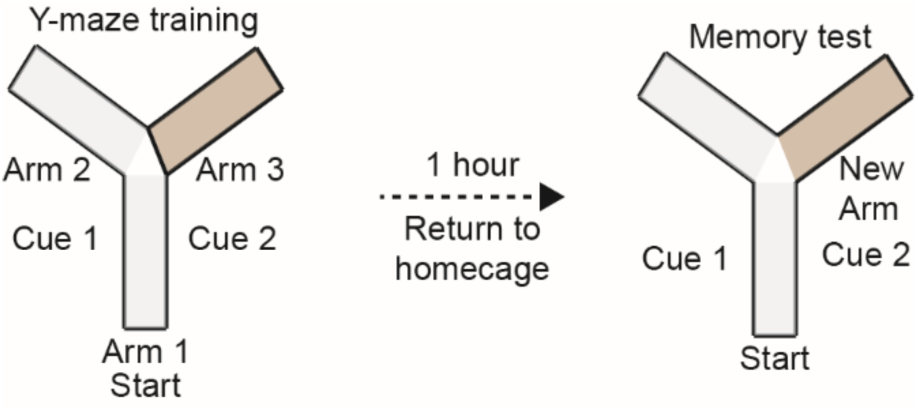
Y-maze spatial memory test. Schematic representation of the modified version of the Y-maze alternation test for assessing short-term spatial memory performance in rodents.

### Supplementary Tables

**Table S1. The differentially expressed proteins upon chronic optogenetic stimulation of the hippocampus in wildtype mice.** Proteins are identified by their gene name together with their UniProt ID number. Average log2 fold differences and P-values have been calculated as described in Methods. The log2 fold change becomes positive when WT+SSFO > WT+tdTom and negative when WT+SSFO < WT+tdTom. P-value determines the statistical significance (*P < 0.05).

**Table S2. Differentially regulated proteins in the hippocampus of 3-month-old 5xFAD.** Gene names and UniProt ID number are used to identify proteins. The log2 fold change becomes positive when 5xFAD+tdTom > WT+tdTom and negative when 5xFAD+tdTom < WT+tdTom. P-value determines the statistical significance (*P < 0.05).

**Table S3. The proteins altered in levels in the hippocampus of 5xFAD mice upon chronic optogenetic stimulation.** Proteins are identified by their gene name and their UniProt ID number is provided. The log2 fold change becomes positive when 5xFAD+SSFO > 5xFAD+tdTom and negative when 5xFAD+SSFO < 5xFAD+tdTom. P-value determines the statistical significance (*P < 0.05).

### Supplementary Data

**Data S1.**
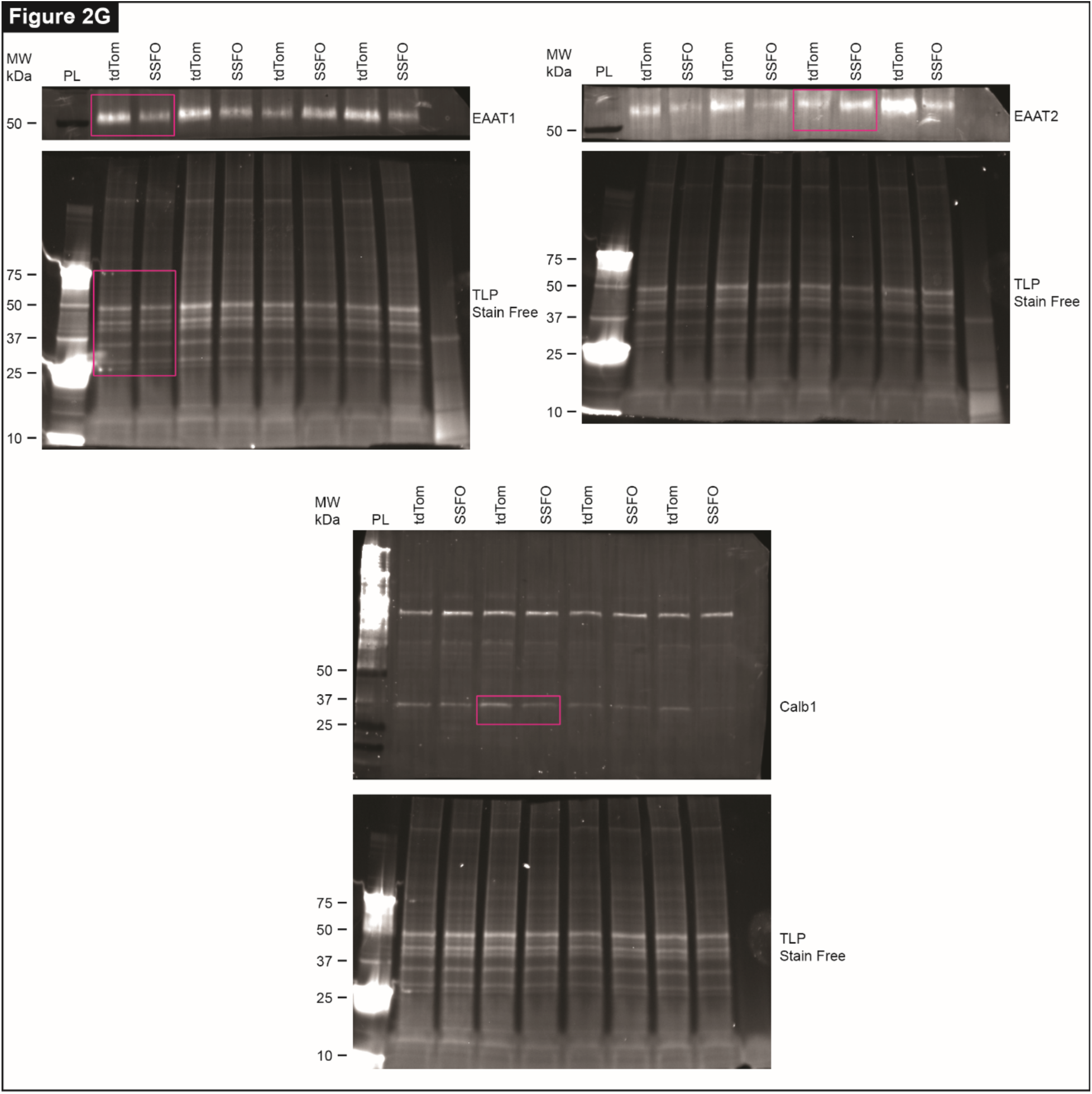
Original, uncropped immunoblots and TGX Stain Free gels related to Fig. 2.

**Data S2.**
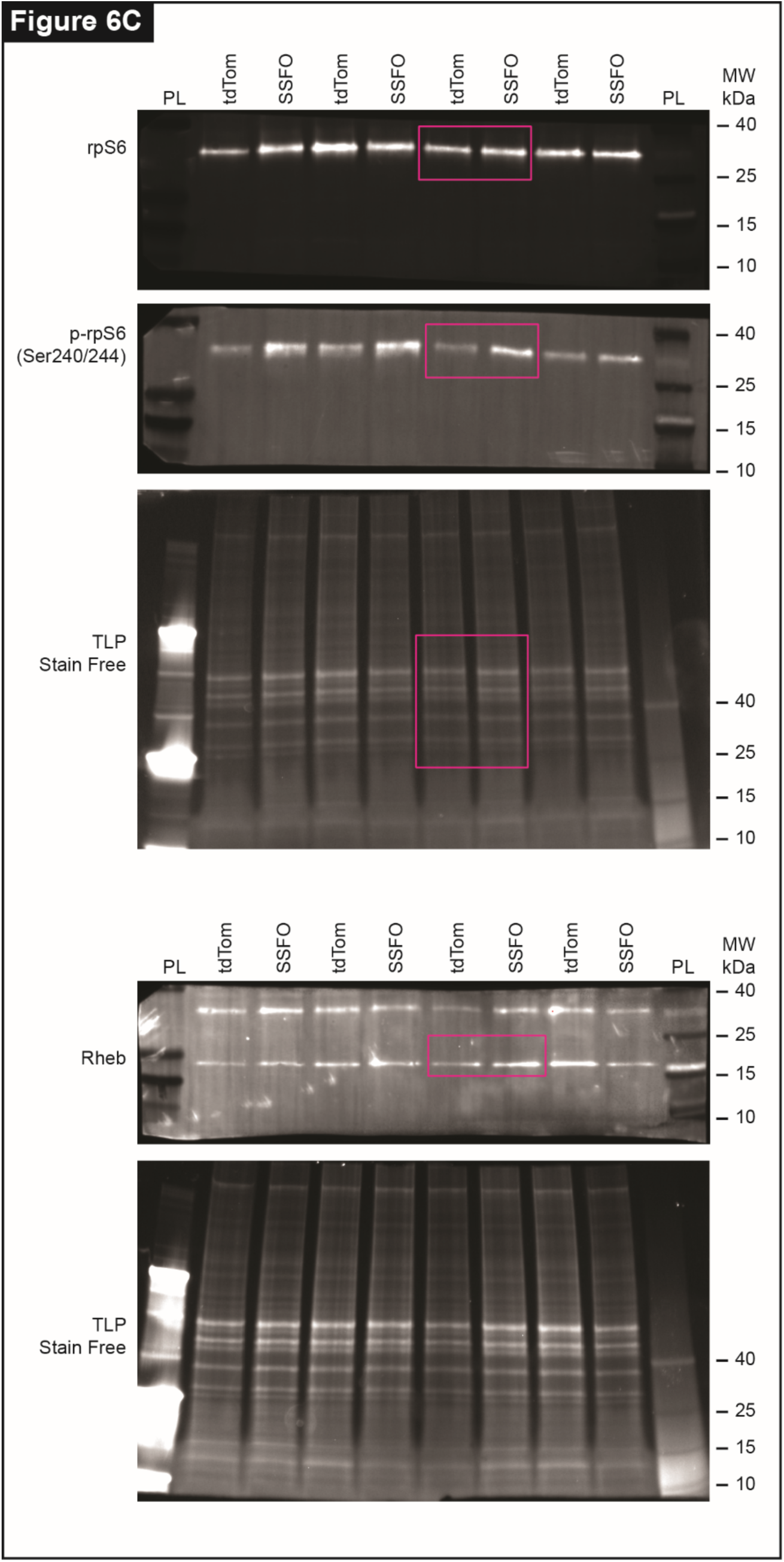
Original, uncropped immunoblots and TGX Stain Free gels related to Fig. 6.

**Data S3.**
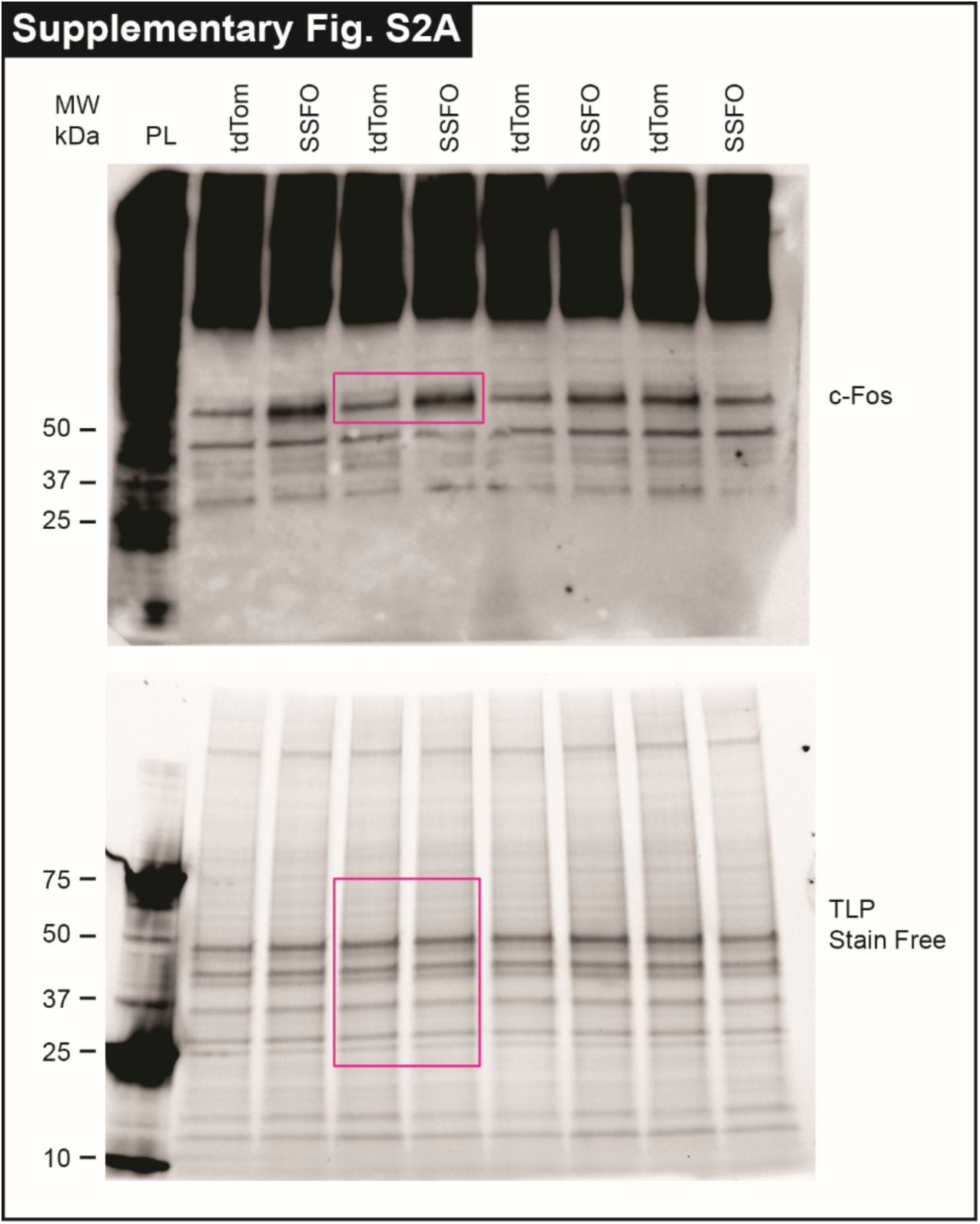
Original, uncropped immunoblots and TGX Stain Free gels related to Fig. S2.

